# Gellan gum-based nutrient medium increases the diversity of cultivated groups of soil microorganisms

**DOI:** 10.1101/2024.11.19.624353

**Authors:** Kulikova Daria Borisovna, Demin Konstantin Alekseevich, Prazdnova Evgenia Valerievna, Mazanko Maria Sergeevna, Kulikov Maksim Pavlovich

## Abstract

Soil microbial communities contain a huge proportion of microorganisms that cannot be cultured on standard microbiological media and are available for study only by molecular methods such as metagenomics. Among them may be found microorganisms-producers of biologically active substances useful for medicine, biotechnology and agricultural activities of humans. Development of approaches for cultivation of such groups may open the way to the creation of fundamentally new antibiotic, antitumor drugs, phyto- and immunostimulants. We have proposed a solid oligotrophic medium for cultivation of soil microorganisms containing gellan gum, which allows to obtain cultures of more rare microorganisms compared to known media. Representatives of phylum Chloroflexota, Deinococcota, Planctomycetota, Actinomycetota, which were absent on agar media, were found in the metagenome of the community grown on this medium

## Introduction

At present, the world scientific community is increasingly addressing the problem of uncultivability of a huge number of large groups of microorganisms. For example, more than 99% of soil microorganism species cannot be cultured by traditional methods [1], and therefore, there is a need to search for alternative ways to obtain cultures of representatives of these groups in laboratory conditions.

Only with the advent of metagenomic sequencing has it become clear how much of the microbial diversity is inaccessible to humans as long as traditional microbiological methods are used. At last count, the GTDB database contains 113104 species of bacteria and archaea, both cultured and uncultured, while the LPSN database contains 20510 species of microorganisms in the category “Names validly published under the ICNP, w/o synonyms”. This category corresponds to species whose isolation in pure cultures is reported in the literature. Thus, a rough estimate of the number of cultured species is 20510/113104, i.e., approximately 18% of the total number of prokaryotes.

Meanwhile, among uncultivated bacteria there may exist species with enormous biotechnological potential. Metagenomic analysis does not yet allow predicting all their properties. Only by combining molecular genetic approaches with fundamentally new cultivation models will it be possible to fully assess the existing biodiversity.

Thus, there is a need to develop media and cultivation methods that will allow us to study microbial consortia in their entirety and significantly expand the list of species cultivated in the laboratory.

Increasing the range of cultivated species can lead to the discovery of new ways of biological control of agricultural pests, increase soil productivity, and can also help in the development of new effective and environmentally safe microbial preparations and biotechnological products.

Various ways to increase the cultivability of soil microbiome representatives have been described in the literature. These include mimicking the composition of soil medium and gas mixture [2, 3], increasing growth time, and reducing the toxicity [4] of media by replacing the gelling agent [5, 6].

In our work, we used the approach of replacing the gelling agent with a less toxic one. Gellan gum was used, which is a very promising gelling agent from the point of view of increasing the cultivability of rare but widespread taxa in soil [7–9].

The aim of this work was to evaluate the taxonomic diversity of cultures grown on media with agar-agar and gellan gum as gelling agents.

## Materials and Methods

The soil used for sowing was selected from the Botanical Garden of the Southern Federal University, and is represented by southern chernozem. The sampling point is represented by the coordinates 47.237541 39.656966.

Two variants of nutrient media were used in the experiment: modified R2A medium diluted 100 times (hereinafter referred to as R2A/100) as oligotrophic medium; Malt extract (hereafter referred to as ME) based medium was used as a copiotrophic medium. The following are the recipes for the nutrient media used in the experiment.

R2A/100 (g/l): glucose – 0.005, K2HPO4 – 0.3, sodium pyruvate – 0.003, peptone – 0.0025, MgSO4 – 0.024, gelling agent – 20.0; ME (g/l): Malt extract – 17.0, peptone– 3.0, gelling agent – 20.0.

All analyses and tests were run in R 4.1.2. All distinctive cultivation experiment variants were conducted in at least 2 biological and at least 6 analytical repetitions. In each case, 2 variants of media were prepared depending on the gelling agent. In the control variants agar-agar was used, in the experimental variants gellan gum was used instead.

Soil dilution (about 10^−4^) was sown on nutrient media, incubation was carried out at t=25°C, CFU counting was carried out on the 10th day. After counting the colonies, the normality of distribution was assessed and the significance of differences between groups was checked by the appropriate statistical test in R software.

### Ecological analysis

The diversity of cultivable microbial community members was assessed as [10] in with minor modifications. Counts of colonies grown during specific day were utilized as distinctive classes for the ecological analysis. In total, 10 classes (for 10 days of incubation) were generated for each experiment condition. The richness and the evenness of the classes were used to express the diversity level. Diversity of a sample was calculated using Shannon diversity index (*H*) formula from vegan package [11]. Evenness of a sample (Pieoou index) was calculated as *H*/ln(*N*), where *N* is the number of classes observed in total during 10 days. Linear model curves were fitted using ggplot2 “geom_smooth()” function with default parameters. Comparison between groups was done using Mann-Whitney test.

### Amplicon data processing

For comparison of gellan gum versus agar gelled R2A/100 medium, biomass from 15 Petri dishes of R2A/100+gellan gum medium and from 15 Petri dishes of R2A/100+agar was rinsed with a physiological solution. For each variant, an equal amount of rinsed biomass was centrifuged, and the pellet was used for DNA extraction. All copies of the 16S rRNA gene were amplified and sequenced. DNA extraction was performed using the FastDNA SPIN Kit for Soil. Metagenomic libraries were prepared following the ‘Preparation of 16S rRNA Gene Amplicons for the Illumina MiSeq System’ protocol. The V3-V4 region of the 16S rRNA gene was amplified using the following primers: forward - TCGTCGGCAGCGTCAGATGTGTATAAGAGACAGCCTACGGGNGGCWGCAG; reverse - GTCTCGTGGGCTCGGAGATGTGTATAAGATACAGGTATCACCHG. Sequencing was carried out on the MiSeq platform (Illumina) with v3 reagents (600 cycles) at the Kazan Federal University Shared Resource Center. Data processing, quality control and operational taxonomic units (OTU) clustering was performed using Mothur [12] software with default parameters.

### Tree construction

To infer closest taxonomic placement of rare taxonomic groups identified in the study, sets of sequences (>800 b.p.) for Chloroflexota and Planctomycetota members were downloaded and combined using RiboGrove [13] and RNAcentral databases. Representative sequences from OTU of interest was aligned with these datasets using mafft (with options “--maxiterate 1000 –localpair” [14]. Trees were inferred using IQ-TREE software with options “-m MF --alrt 1000” [3] and visualized using iTol web service [15].

## Results

The results obtained during the experimental part are presented below.

As can be seen in Figure 1, the use of gellan gum as a gelling agent in combination with oligotrophic nutrient medium allows to achieve a more significant increase in CFU at plating of the same material in comparison with media where the gelling agent is agar-agar. In addition, this effect is maintained on nutrient medium ME with a high concentration of nutrients. A significantly lower number of CFU was also observed on “starved” nutrient media than on the corresponding variants with low nutrient concentration (R2A, SA)It can be observed that Shannon’s index and Pielou’s index are higher on the medium where gellan gum was used as gelling agent. In addition, on the oligotrophic nutrient medium R2A/100, the difference of these two parameters is statistically significant. It follows that in the gellan gum culture medium there is a greater diversity of unique, but phylogenetically close taxa, which are more uniformly represented than taxa in the agar-agar-based medium culture medium. The Shannon Index characterizes community biodiversity, while the Pielou Index demonstrates the equal representation of species within a community.

**Figure 1.**
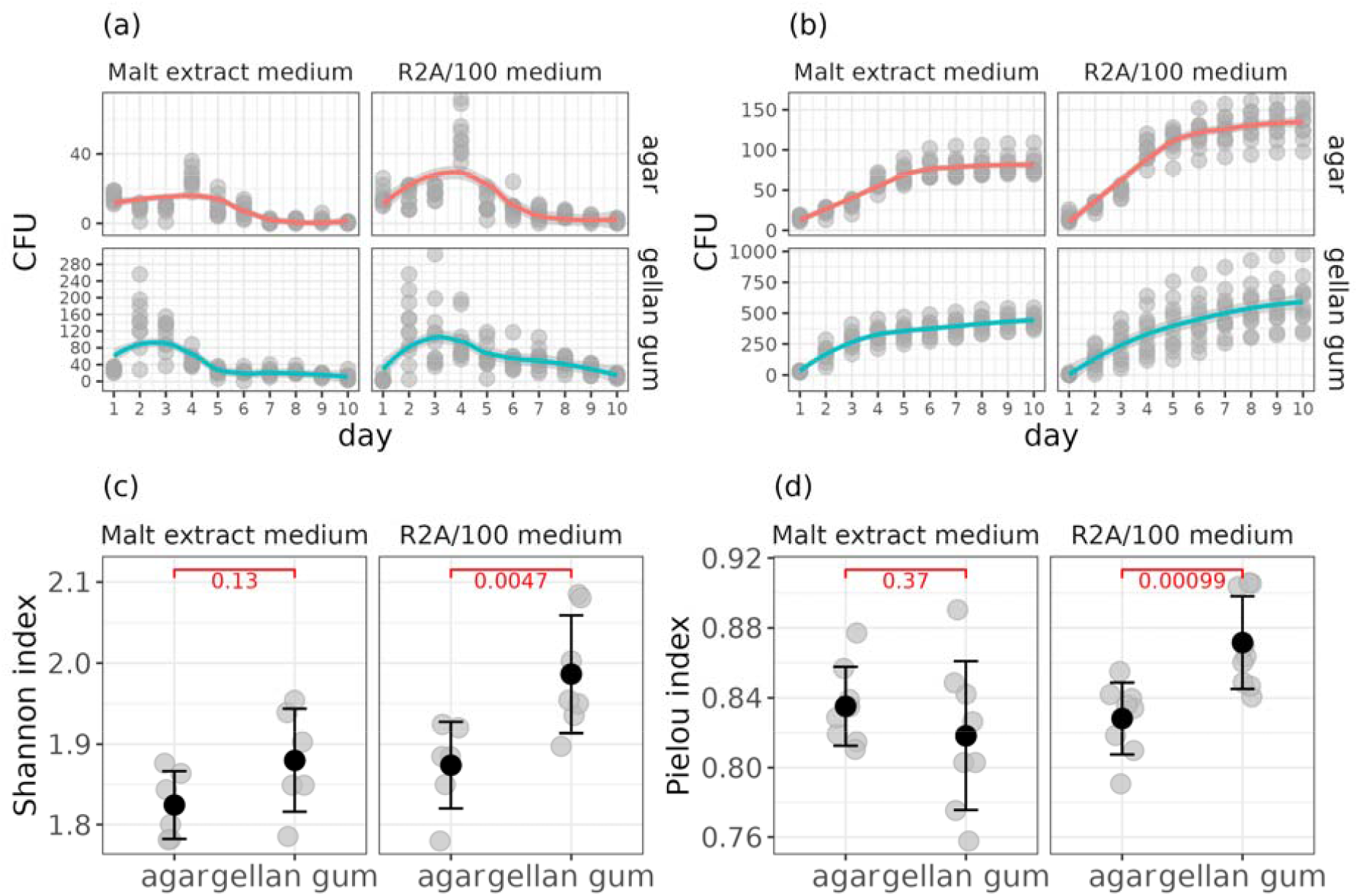
(a) – CFU count on different media by day; (b) – cumulative sums of CFU counts during the incubation period; (c) – Mann-Whitney test comparison of Shannon’ diversity indexes calculated based on the abundance of ten classes; (d) – Mann-Whitney test comparison of Pielou’ evenness indexes calculated based on the abundance of ten classes.

As shown in Figure 2, analysis of the prokaryote families unique to the two experiments indicates that the gum culture medium is dominated by microorganisms phylogenetically distant from the typical cultured taxa. Families unique to the agar-agar culturerome belong to phyla that are most often isolated from soil in laboratory conditions: Pseudomonadota, Actinomycetota and Bacteroidota. The phylum Chloroflexota and Planctomycetota, which are widely distributed in soils but rarely cultivated in laboratory conditions, were observed in the variant with gum. The highest number was observed for the family JG30-KF-CM45, representatives of which were not observed in pure cultures. Representatives of the family Isosphaeraceae, also widely represented in soils on agar-agar media, were not detected in this work. Finally, representatives of the order Solirubrobacterales were previously isolated both on oligotrophic media and on media with the addition of superoxide dismutase [16].

**Figure 2.**
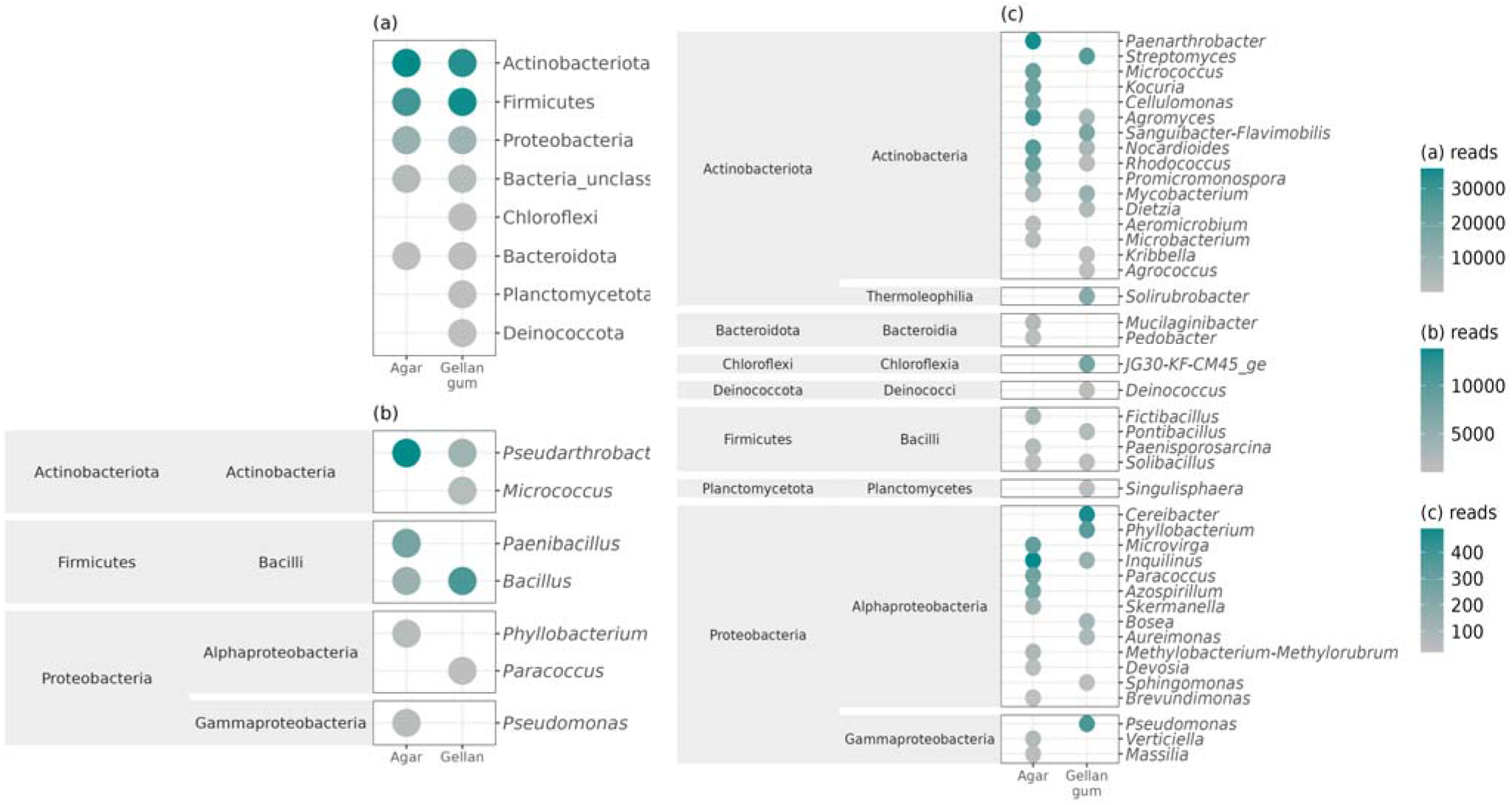
(a) – abundance of bacterial phyla grown on R2A/100 medium with agar and gellan gum as gelling agents; (b) – abundance and taxonomy of dominant bacterial genera grown on R2A/100 medium with agar and gellan gum as gelling agents; (c) – abundance and taxonomy of minor bacterial genera grown on R2A/100 medium with agar and gellan gum as gelling agents

Figure 3 demonstrates, that for Planctomycetota affiliated amplicons, the closest tree placement was Singulisphaera acidiphila. In contrast, for Chloroflexota amplicons we were unable to resolve the closest tree placement at the species level. We conclude that these amplicons derive from yet unknown member of Sphaerobacteraceae family, which, as it seems, was grown in our experiment using gellan gum-gelled R2A/100 medium.

**Figure 3.**
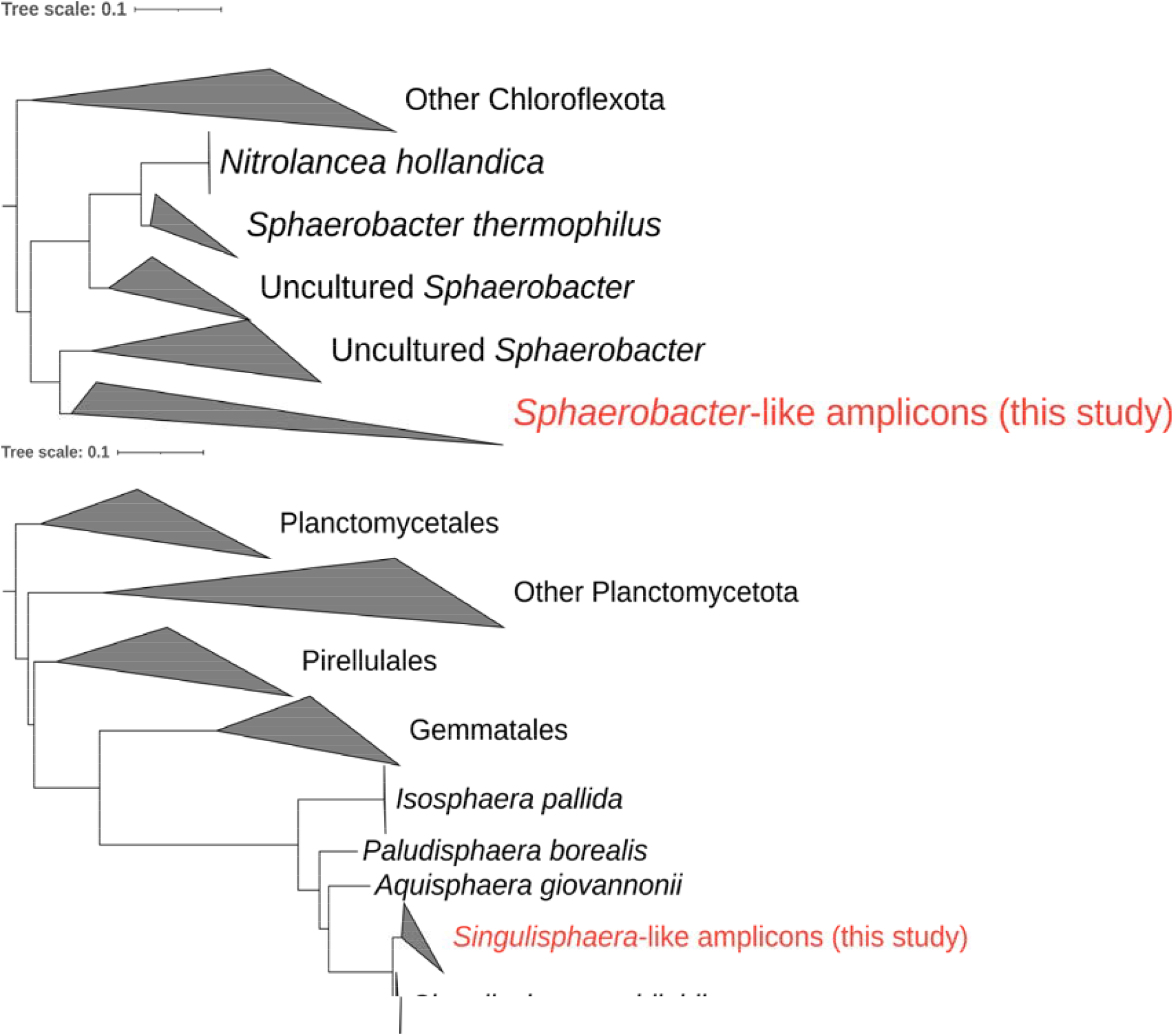
Phylogenetic trees showing the position of Chloroflexota (on the top) and Planctomycetota (at the bottom) amplicons relative to the different taxonomic groups of the corresponding phyla.

## Discussion

The phenomenon of microbial uncultivability may be caused by unfavorable abiotic conditions [1, 17] such as suboptimal temperature values, nutrient deficiencies, elevated salt concentrations [18], pH fluctuations [19], osmotic stress, reactive oxygen species, and heavy metal exposure [20]. It is hypothesized that many microorganisms that do not grow on rich nutrient media belong to K-strategists. Such bacteria grow slowly and require a different ratio of nutrients to biomass than fast-growing r-strategists [21, 22]. It may be difficult to recreate conditions that are as close as possible to the optimum zone of unculturable groups of microorganisms in laboratory conditions. In addition, a significant part of unculturable representatives require extreme growth conditions, in particular, maintenance of an anaerobic atmosphere, high temperature values, and elevated salt concentrations

Biotic interactions with representatives of other groups can also have a significant impact on the cultivability of some bacterial phyla [23]. For example, these interactions can be both positive [24] and negative [25]. In the first case, microorganisms are capable of synthesizing various growth factors, vitamins, and amino acids, which is a vital condition for those groups lacking key genes for synthesizing these compounds. In the case of negative biotic influence of some groups of bacteria on others, this refers mostly to competition for nutrients and growth factors, as well as to inhibitory influence by synthesizing secondary metabolites.

The existing approaches to increasing the number of cultivated groups in the study of soil microbiota can be summarized in two principal directions:

1. Increasing the number of representatives of such groups, which under normal conditions are largely displaced by fast-growing species. As a rule, such groups represent an aboriginal part of the soil microbial community.
2. The isolation of hard-to-cultivate species and species that have never been cultured in the laboratory before.

The first approach is most commonly used to:

1. mimicking the composition of the soil medium. The extraction of soil extract by non-aqueous solvents is also promising [3].
2. simulation of the gas mixture characteristic of soils. Numerous literature data on the composition of the gas mixture favorable for the cultivation of soil microbiota suggest that the carbon dioxide concentration is the determining factor [2].
3. The “comfort time” strategy [4]. Long-term incubation approach with periodic replacement of the carbon source. Based on the observation that many soil bacterial species have long generation times [26, 27] and proliferate over a narrow range of environmental conditions, especially a range of nutrient concentrations.

In the second approach, researchers are more likely to focus on:

1. Reducing the toxicity of the nutrient media base. The approach is based on the use of non-agar gelling agents (carrageenan, gelrite, phytagel) and the addition of antioxidants and reactive oxygen species quenching enzymes to the medium. It has been shown that both agar-agar itself and the reactive oxygen species formed in the agar gel can reduce the activity or completely inhibit the growth of certain groups of prokaryotes [28]. The use of other gelling agents has previously achieved the cultivation of rare bacteria and a more efficient cultivation procedure in general [5, 6]. Presumably, for many bacteria living in natural nutrient-poor environments such as soil, seawater, and freshwater, the presence of large amounts of nutrients in culture media may negatively affect growth and development rates [21, 22]. Therefore, repeated dilution of the nutrient medium resulted in the growth of unculturable taxa on it [29, 30].
2. Selective growth suppression. The approach is based on the possibility of using a large variety of substances in the nutrient medium that selectively suppress the growth and reproduction of typically isolated groups. It has been shown, for example, that the addition of chromate ions to the medium gives a selective advantage to mycelial actinobacteria [31]. In addition, this effect can be prevented by using the dilution-to-extinction method in order to reduce the concentration of cells in the inoculum and isolate representatives of the numerically most represented groups, which, as a rule, belong to the autochthonous part of the microbocenosis [32].
3. Application of the ichip method [33]. The method is based on placing cultured bacteria in their natural habitat, which contains the necessary composition of substances and molecular signals.

At the moment, there is a great variety of nutrient bases for isolation of microorganisms from soils. To increase the diversity and CFU number of the cultured part of the soil microbiome, it is often resorted to changing the composition of nutrient media components or modifying the cultivation technology. One of the most significant components affecting the cultivability of soil microorganisms is the gelling agent in the medium.

The most common gelling agent in laboratory microbiological practice is agar-agar, a polysaccharide derived from red algae of the genera Phyllophora, Gracilaria, Gelidium, Ceramium. Agar-agar has a melting point of 85°C, it is transparent, has optimal diffusion characteristics, and is also not susceptible to enzymatic cleavage by most microorganisms; however, some marine [34, 35] and soil microorganisms are able to utilize it for their needs. Due to these qualities, at this point in time, agar has such a wide distribution. In the vast majority of cases, agar-agar is used because of its low cost, relative biological inertness and ease of preparation of solid media.

Despite this, agar-agar has disadvantages such as the presence of impurity compounds that reduce the transparency of the gel formed, as well as toxicity to some groups of microorganisms. It has been observed that most of the naturally occurring microorganisms cannot grow on agar-agar media. In particular, the interaction between agar and phosphate during the preparation of nutrient media leads to the formation of hydrogen peroxide (H_2_O_2_), which negatively affects the ability to cultivate bacteria on the medium [36].There is also evidence that traces of furan-2-carboxylic acids have been found in commercial brands of agar, which may also be the cause of growth inhibition of rare groups of microorganisms [9].

Since it is unknown how many taxonomic or physiological groups of prokaryotes are unable to grow in the presence of agar-agar, the use of other gel-forming polysaccharides, such as guar gum, xanthan gum, gellan gum, carrageenan, silica gel, and izubgol, has been actively discussed [36].

One of the most promising substitutes for agar-agar is gellan gum, which is a mixture of oligosaccharides produced by some species of bacteria genus Sphingomonas. By molecular structure it is a linear heteropolysaccharide consisting of repeating links of beta-D-glucose, beta-D-glucuronic acid, beta-D-glucose and alpha-L-rhamnose. Gellan gum forms dense transparent thermostable gels, which allows the cultivation of thermophilic prokaryote species due to its high melting point (∼110°C). Gellan gum gels are also stable in a wide pH range [37].

As our study showed, the use of gellan gum as a gelling agent instead of agar-agar allows to achieve a more significant increase in CFU when sowing the same material on different types of media. The effect remained stable on oligotrophic media, on nutrient-rich media, and even on starvation media. Significantly lower CFU counts were also observed on “starved” nutrient media than on the corresponding variants with low nutrient concentrations (R2A, SA). This allows us to exclude the possibility that this effect is caused by the use of oligosaccharides in gellan gum as a carbon source by microorganisms.

The greatest difference in the number of CFU was on R2A/100 medium, so the cultures grown on this medium were sequenced by 16S-RNA. Sequencing showed that not only the number of CFU, but also the structure of the grown cultures differed. In the gellan gum variant, on the one hand, the number of classified families was lower, but, on the other hand, the number of unique genera, including unclassified respects, was higher. That is, replacing agar with gellan gum does not simply increase the number of bacteria grown, but affects the diversity of the culturomere as a whole (Figure 2). By changing the gelling agent, we must realize that we will gain some of the families not cultured on agar, but we will lose some of the families not cultured on gum, such as those belonging to phylum such as Pseudomonadota, Actinomycetota and Bacteroidota. According to the Shannon index, the agar-agar culture medium is generally more diverse than the gellan gum culture medium. On the other hand, good growth of phylum Chloroflexota and Planctomycetota, the growth of which is difficult to achieve on agar agar, as well as family JG30-KF-CM45, representatives of which were not observed in pure cultures, was observed on gellan gum. There is evidence that the representation of this family is associated with the activity of soil enzymes [38] and organic carbon accumulation [39].

Based on the results obtained during the study, we can conclude that the bacterial communities grown on media where gellan gum was used as a gelling agent had greater biodiversity, which allows us to formulate the conclusion that the use of gellan gum as a gelling agent instead of agar allows the growth of more diverse microorganisms, including those from rarely seeded taxonomic groups. This phenomenon may be due to the absence of inhibitory substances in the composition of agar media.

The literature describes cases of cultivation of such rare taxa as Acidobacteria, Chloroflexi, Gemmatimonadetes, Planctomycetes, and Verrucomicrobia on gellan gum. In particular, in the work of Kagomata and Tamaki, the number of colony forming units (CFU) was 10 times higher on the medium where gellan gum was used as gelling agent instead of agar-agar [37]. In a study by Tamaki and Hanada on gellan gum, 108 isolates were isolated, 50 of which showed high phylogenetic novelty. These isolates belonged to the types Proteobacteria, Firmicutes, Bacteroidetes and Verrucomicrobia, with 18 isolates having very low sequence similarity (<94%) to the closest species in the available 16S-rRNA gene sequence database [40]. Also in a study by Janssen et al. on gellan gum medium, they were able to identify isolates affiliated with major bacterial groups such as Proteobacteria, Actinobacteria, Acidobacteria, and Verrucomicrobia. Based on the similarity of 16S rRNA gene sequences, some of these isolates belonged to bacterial lines that had not been isolated in pure cultures before [41].

Thus, the use of modified R2A medium with gellan gum as a gelling agent will make it possible to isolate from soil microorganisms previously considered unculturable, and thus significantly expand the area of screening of bacteria with the necessary functions. The resulting microorganisms may possess qualities not represented in known groups of soil microorganisms, or significantly m^4^ore pronounced. The nutrient medium is suitable for the search of new groups of microorganisms for use in various fields of economic activity, in particular for the search of microorganisms with antimicrobial activity, development of biologically safe methods of control, organic fertilizers, search of microorganisms capable of entering into symbiotic relations with economically important plants, fungicides, plant growth regulators, antibiotics and other applications.

## Conclusion

In this study, we showed significant differences in the composition of microorganisms cultured on agar-agar and gellan gum media. On the oligotrophic medium, which is a modified R2A medium, the biodiversity of cultures grown on gellan gum was higher compared to the biodiversity of cultures grown on agar-agar. In addition, bacteria rarely or not at all capable of growing on agar-agar media were found on gellan gum medium. Thus, the use of oligotrophic media with gellan gum may increase the number of cultured microorganisms available for further study and application.

The result of the work is protected by the Russian Federation Patent № RU 2 818 084 C1

The study was carried out in the Laboratory of Molecular Genetics of Microbial Consortia of the Southern Federal University, funded by the Strategic Academic Leadership Program of the Southern Federal University (“Priority 2030”).

